# Cancer stemness, intratumoral heterogeneity, and immune response across cancers

**DOI:** 10.1101/352559

**Authors:** Alex Miranda, Phineas T Hamilton, Allen W Zhang, Etienne Becht, Artur Mezheyeuski, Jarle Bruun, Patrick Micke, Aurélien De Reynies, Brad H Nelson

## Abstract

Regulatory programs that control the function of stem cells are active in cancer and confer properties that promote progression and therapy resistance. However, the impact of a stem cell-like tumor phenotype (“stemness”) on the immunological properties of cancer has not been systematically explored. Using gene expression-based metrics, we evaluate the association of stemness with immune cell infiltration and genomic, transcriptomic, and clinical parameters across 21 solid cancers. We find pervasive negative associations between cancer stemness and anticancer immunity. This occurs despite high stemness cancers exhibiting increased mutation load, cancer-testis antigen expression, and intratumoral heterogeneity. Stemness was also strongly associated with cell-intrinsic suppression of endogenous retroviral expression and type I interferon signaling and increased expression of several therapeutically accessible signaling pathways. Thus, stemness is not only a fundamental process in cancer progression but may represent a unifying mechanism linking antigenicity, intratumoral heterogeneity, and immune suppression across cancers.

## Introduction

Immunotherapy has recently emerged as an important therapeutic modality for a broad range of cancers. In particular, major therapeutic gains have been made using antibodies against the CTLA-4 and PD-1 pathways, as well as adoptive cell therapies using natural or engineered T cells (June et al., 2018; Ribas and Wolchok, 2018). Despite these advances, however, the reality remains that today’s immunotherapies are effective for only a minority of patients. Among several challenges facing the field, a large proportion of solid cancers are non-permissive to lymphocyte infiltration (immunologically “cold”), protecting them from cytolytic attack by lymphocytes such as CD8+ T cells (Gajewski, 2015). With improved understanding of the mechanisms underlying the cold tumor phenotype, the benefits of immunotherapy could potentially be extended to a much larger number of patients.

Mounting evidence suggests that, like normal tissues, tumors can possess a hierarchical structure of phenotypically diverse cell populations with varying capacities for self-renewal and differentiation. The cancer stem cell (CSC) hypothesis posits that a subpopulation of cells resides at the top of the cellular hierarchy and sustains the long-term maintenance of neoplasms (Balttle and Clevers, 2017). This hypothesis provides compelling explanations for clinical observations such as therapeutic resistance, tumor dormancy, and metastasis (Nassar and Blanpain, 2016). CSCs have been identified in a variety of human tumors, as assayed by their ability to initiate tumor growth in immunocompromised mice (Al-Hajj et al., 2003; O’Brien et al., 2007; Singh et al., 2004). While considerable controversy remains as to how best to define CSCs and the extent to which different tumor types exhibit this hierarchical organization, it is increasingly clear that stem cell-associated molecular features, often referred to as ‘stemness’, are biologically important in cancer (Kreso and Dick, 2014). Indeed, a negative association between stemness and prognosis has been reported for a wide variety of cancers (Liu et al., 2007; Merlos-Suarez et al., 2011; Ng et al., 2016). Moreover, stemness appears to be a convergent phenotype in cancer evolution (Chen and He, 2016; Greaves, 2013), suggesting it is a fundamentally important property of malignancy.

The evolution of transformed cells in the tumor microenvironment is shaped by diverse selective pressures, including the host immune response. Experimental work has shown that embryonic, mesenchymal, and induced pluripotent stem cells possess immune modulatory properties, while resistance to immune-mediated destruction has also recently been shown to be an intrinsic property of quiescent adult tissue stem cells (Agudo et al., 2018). These properties can also be shared by CSCs (Bruttel and Wischhusen, 2014; Maccalli et al., 2014). In cancer, immune selection has been shown to drive tumor evolution toward a high-stemness phenotype that inhibits cytotoxic T cell responses (Noh et al., 2012). Indeed, a recent analysis of the Cancer Genome Atlas (TCGA) revealed negative associations between stemness and some metrics of tumor leukocyte infiltration (Malta et al., 2018). Stemness has also been proposed as a driver of intratumoral heterogeneity, with CSCs proposed as the unit of selection in cancer (Greaves, 2013). Consistent with this, we (Zhang et al., 2018) and others (Safonov et al., 2017) have reported negative associations between immune cell infiltration and intratumoral heterogeneity.

Motivated by these observations, we hypothesized that a stemness phenotype of cancer cells may confer immunosuppressive properties on tumors, resulting in immunologically cold microenvironments that both foster and maintain intratumoral heterogeneity. To address this, we performed an integrated analysis of stemness, immune response, and intratumoral heterogeneity across cancers. We recover pervasive negative associations between antitumor immunity and stemness, and strong positive associations between stemness and intratumoral heterogeneity. We further find that cancer cell lines with high stemness have cell-intrinsic immunosuppressive features, suggesting that immunologically cold microenvironments can arise due to the presence of high-stemness cancer cells. We propose that cancer stemness provides a unifying mechanism that links tumor antigenicity, intratumoral heterogeneity, immune suppression and the resulting evolutionary trajectories in human cancer.

## Results

### Derivation and comparison of stemness signatures

We inferred tumor stemness from cancer transcriptomes using single sample gene set enrichment analysis (ssGSEA) with a modified version of a gene set developed by Palmer and colleagues to measure the level of plasticity and differentiation of mesenchymal stem cells, pluripotent stem cells, terminally differentiated tissues, and human tumors across >3200 microarray samples (Palmer et al., 2012) (See methods). Intriguingly, the authors of this gene set identified a cluster of ‘immune’ genes that negatively loaded the principal components they used to infer stemness; however, they did not further explore this relationship. To adopt this gene set for use in ssGSEA and avoid biasing our analysis towards recovering negative associations between stemness and immunity, we omitted this immune gene cluster from our signature. We also omitted cell proliferation markers to avoid recovering a signature of proliferation rather than stemness (Ben-Porath et al., 2008). The resulting 109-gene stemness gene set (**Table S1A**) was applied to RNA sequencing data from 8,290 primary cancers from 21 solid cancer types in TCGA, omitting instances in which multiple tumor samples were acquired from a single patient.

When applied to TCGA samples, our stemness signature showed only a moderate correlation with a recently published mRNA-based stemness signature (‘mRNAsi’; N = 7,429 overlapping samples; Spearman’s ρ = 0.43; **Figure S1A**) that was derived using a novel one class logistic regression (OCLR) (Malta et al., 2018). Despite these differences, when evaluating our stemness signature against the validation dataset of Malta et al. (GSE30652), the two signatures yielded correct classification for 238/239 samples; the one exception was a parthenogenetic stem cell sample that scored below the highest non-stem cell sample (**Figure S1B and S1C**). When applied as a classifier to all tissue classes and cell types in this dataset, our signature showed comparable performance to that of Malta et al. (multiclass AUC 0.92 vs. 0.91, respectively) (See methods). Thus, our signature provides a useful metric of stemness that is largely concordant with the mRNAsi signature of Malta et al. when applied to the above dataset but, as described below, yields contrasting results when applied to human cancers.

### Stemness varies across cancers and predicts patient survival

Using our signature, we found that stemness varied strongly across TCGA samples, with cancer type explaining 54% of the variation (ANOVA; adjusted R^2^; **Figure 1A**). Consistent with prior reports of stemness being a negative prognostic factor (Liu et al., 2007; Merlos-Suarez et al., 2011; Ng et al., 2016), we found a strong negative relationship between median stemness and median overall survival across cancer types (**Figure 1B**; ρ = −0.60; P = 0.004; N = 21 cancers). Within cancer types, univariate Cox regression likewise showed stemness to be significantly negatively prognostic for overall survival in the majority of cancers (Cox proportional hazards; P < 0.05), and significantly positively prognostic for none (**Figure 1C**) (See methods), underscoring the relevance of this signature both within and across cancers. We also noted a significant decrease in the magnitude of the hazard associated with stemness within cancers as median sternness increased (ρ = −0.55; P < 0.01), pointing to a potential threshold effect with a saturating hazard in cancers with higher average stemness. For reference, we compared the prognostic association of our ssGSEA-based stemness with the OCLR-based mRNAsi, for which positive associations in some cancers were reported (Malta et al., 2018). Using pan-cancer Cox regressions stratified by cancer, we found ssGSEA-based stemness to be substantially more predictive of survival in this linear modelling framework (log hazard ratio = 0.23 ± 0.03 (coefficient ± SE); P < 10^−15^ vs. not significant for mRNAsi), demonstrating our signature more effectively uncovers the negative outcomes associated with high-stemness cancers expected from previous reports.

**Figure 1.**
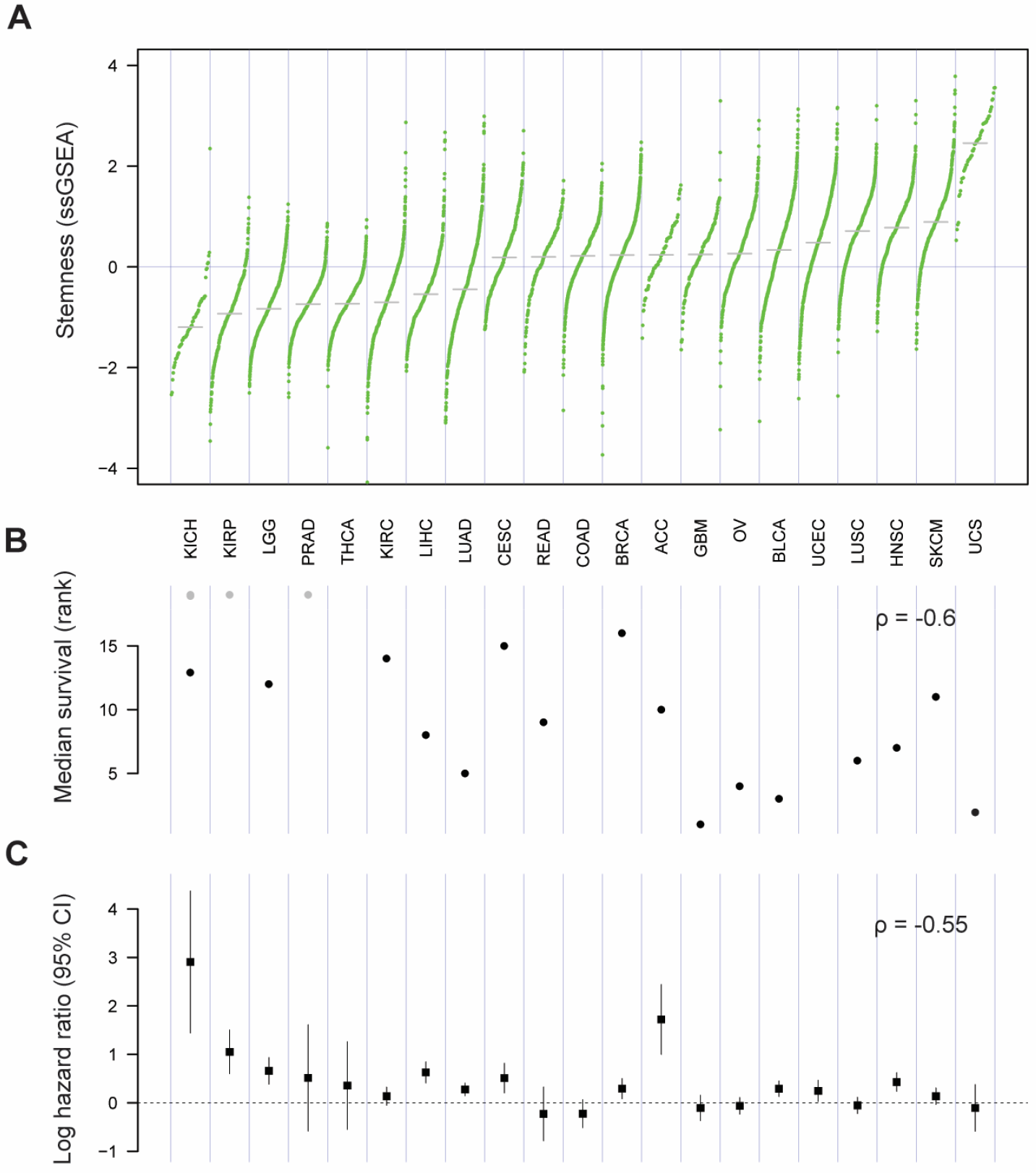
Stemness and survival across cancers. **A)** Stemness score varies widely across 21 solid cancers from TCGA. Each point represents an individual case, and cancer types are ordered by median stemness score (z-scored ssGSEA). **B)** Median survival decreases with increasing median stemness (P = 0.004). Gray points represent cancers in which median overall survival times were not evaluable. **C)** Stemness associates with poor outcome within cancers. Log hazard ratio (± 95% CI) for the association of stemness with overall survival is shown. Hazard decreases with increasing average stemness of cancers (P = 0.008). See also **Table S1A**.

### Stemness negatively associates with immune cell infiltration across solid cancers

To evaluate the relationship between stemness and antitumor immunity, we generated signatures of predicted immune cell infiltration for each patient sample using xCell, an ssGSEA-based tool that infers cellular content in the tumor microenvironment (Aran et al., 2017) (See methods). CD8+ T cells, which have a well-established association with favorable prognosis in a majority of solid cancers (Gooden et al., 2011), showed a clear negative association with stemness for most cancers (**Figure 2A**). We also considered other cell types important for antitumor immunity, including NK cells (Larsen et al., 2014) and B cells (Li et al., 2016; Nelson, 2010), and again observed recurrent negative associations with stemness (**Figure 2A**). Additional cell types such as CD4+ T cells, T regs, and neutrophils showed more variable associations with stemness, indicating this relationship does not apply to all infiltrating immune cell populations in all cancers (**Figure 2A**).

**Figure 2.**
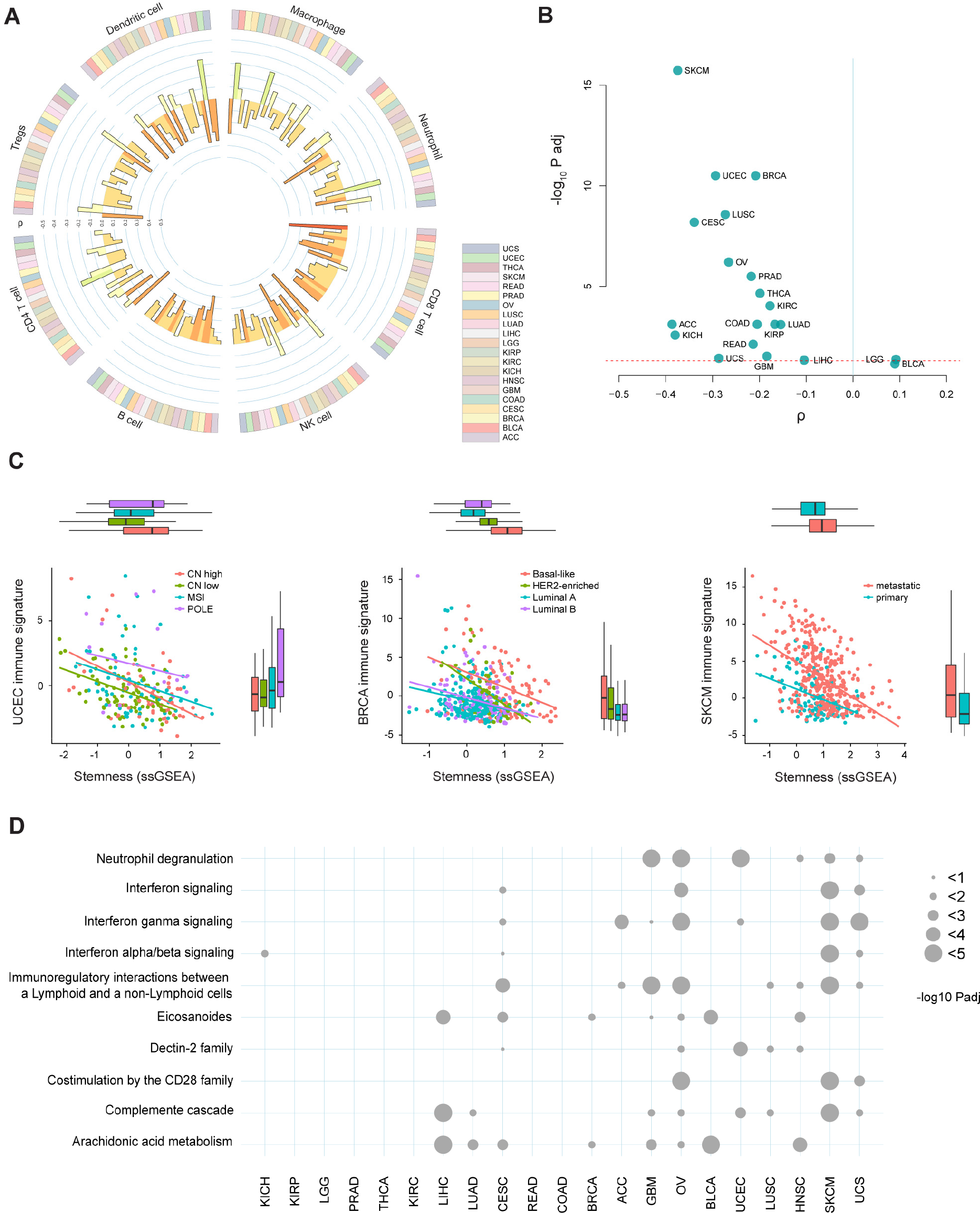
Stemness negatively associates with immune cell signatures. **A)** Circos plot showing the association between stemness score and the presence of 8 inferred immune cell types across 21 cancer types (colored bars in outer ring). The color and height of the inner bars represent the Spearman correlation ρ values for each cell type and cancer type. **B)** Volcano plot showing the association between stemness score and immune signature (sum of z-scored signatures of CD8+ T cells, NK cells, and B cells) for each cancer. The dashed line indicates P_adj_ = 0.05. **C)** Association between stemness score and immune signature in the different molecular subtypes of endometrial (UCEC) and breast (BRCA) cancer, and within primary and metastatic melanoma (SKCM) samples (P_adj_ < 10^−7^). Each point represents one case. Colors indicate the different molecule subtypes of UCEC and BRCA, or sample types for SKCM. CN = copy number, MSI = microsatellite instable, POLE = polymerase epsilon. **D)** Reactome pathway enrichment analysis of the top 1,000 genes up-regulated in low stemness (<20^th^ percentile) vs. high stemness (>80^th^ percentile) samples. The size of each point reflects −log_10_ adjusted P values. See also **Figure S2,3,4** and **Table S1**.

To generate a robust single score for antitumor immunity (hereafter referred to as ‘immune signature’), we aggregated xCell predictions for CD8+ T cell, NK cell, and B cell infiltration, reasoning that these cells represent important anticancer effector cells across diverse cancers (Melero et al., 2014; Sarvaria et al., 2017). Indeed, this immune signature was significantly associated with increased survival in the majority of cancers (**Figure S2A**; overall survival curves for patients stratified by median stemness and immune signature are shown in **Figure S2B**). A notable exception was certain kidney cancers, where negative associations were observed, in accord with prior reports for these malignancies (Jochems and Schlom, 2011). Consistent with our analyses using single immune cell type scores (**Figure 2A**), this immune signature showed a negative association with stemness within nearly all cancers (**Figure 2B**). This negative association was also recovered using other published stemness gene sets (Ben-Porath et al., 2008; Bhattacharya et al., 2004; Shats et al., 2011) (**Table S1B**), most clearly with stemness scores reflecting NANOG, SOX2 and MYC signaling, and to a lesser extent with those reflecting embryonic stem cell programs (**Figure S3**). Furthermore, when we used recently published CIBERSORT infiltration scores (Thorsson et al., 2018) in place of our immune signature, we found that increased stemness was associated with strong polarization of infiltrating leukocyte populations towards a macrophage-dominant, CD8+ T cell-depleted composition for most cancers (linear model controlling for cancer type; P < 10^−15^ for both cell fractions).

Whereas the preceding analyses were performed *within* cancer types, we also evaluated the relationship between stemness and immune signature *across* cancer types. Unexpectedly, we found no significant association between median immune signature and median stemness score across cancers (P = 0.82). This suggests that factors other than stemness control the differences in immune cell infiltration across cancer types, while the association with stemness applies within individual cancer types. It also demonstrates that our stemness metric is not recovering negative associations with immunity simply due to the lower tumor purity that is inextricably associated with the presence of infiltrating immune cells.

We next examined whether the negative association between immune signature and stemness was influenced by tumor subtype or stage. Here, we focused on breast and endometrial cancers, which have well-characterized subtypes with strong prognostic associations, and melanoma, for which both primary and metastatic samples are available within TCGA. In breast cancer, stemness varied markedly across subtypes (ANOVA; adjusted R^2^ = 0.26), with the basal subtype having the highest stemness, as expected (Ben-Porath et al., 2008), and the luminal-A subtype the lowest (**Figure 2C**). In endometrial cancer, the highest stemness score was observed in high-copy number alteration (CN-high) and polymerase-epsilon mutant (POLE) tumors, and the lowest stemness score was seen in low-copy number alteration tumors (CN-low) (Tukey HSD; P = 0.003; CN-high vs. CN-low tumors). Finally, we observed a substantially higher stemness score in metastatic compared to primary melanoma lesions (**Figure 2C**). In all these cancers, we observed recurrent negative associations between stemness and immune signature which remained significant when controlling for cancer subtype and tumor purity (see below; linear models; P < 10^−7^).

To investigate in an unbiased manner whether processes apart from antitumor immunity negatively correlate with stemness, we conducted differential expression tests to identify gene expression patterns associated with the lowest versus highest stemness quantiles (< 20^th^ versus > 80^th^ percentiles) for each analyzed cancer type (See methods). Even with this unbiased approach, nearly all the pathways recurrently enriched in low-stemness samples within a cancer were immune-related (**Figure 2D**). Recognizing that the presence of non-malignant cells (e.g. immune, stromal, endothelial cells) can confound expression analyses of bulk-sequenced tumor samples by diluting tumor-specific expression signatures, we performed additional analyses to control for such effects. First, using recently published estimates of purity across TCGA (Aran et al., 2015), we found that the association between tumor purity and stemness was negligible and failed to reach significance in a pan-cancer linear model controlling for cancer type (P = 0.23). Second, we re-fit the differential expression models described above to control for tumor purity and repeated the pathway enrichment analyses, still finding that immune-associated pathways were enriched in low-stemness tumors (**Figure S4**) (See methods).

### Stemness associates with immunologically cold cancers measured via IHC

To confirm the negative association between stemness and lymphocyte infiltration, we turned to three patient cohorts with matched immunohistochemistry (IHC)-based T cell infiltration scores and gene expression data suitable for computing a stemness score (See methods). Using a cohort of 33 colorectal cancer patients (Becht et al., 2016), we found a strong negative association between stemness and total infiltrating CD3+ T cells (ρ = −0.63; P < 0.001; **Figure 3A**). Furthermore, in this cohort the xCell-based immune signature was strongly correlated with infiltrating CD3+ cells (sρ = 0.69; P < 0.001), supporting its fidelity for measuring immune-cell infiltration. With this validation in hand, we compared the xCell-based immune signature and stemness for the total patient cohort with available microarray data (N = 585), and again observed a clear negative correlation (ρ = −0.22; P < 10^−7^).

**Figure 3.**
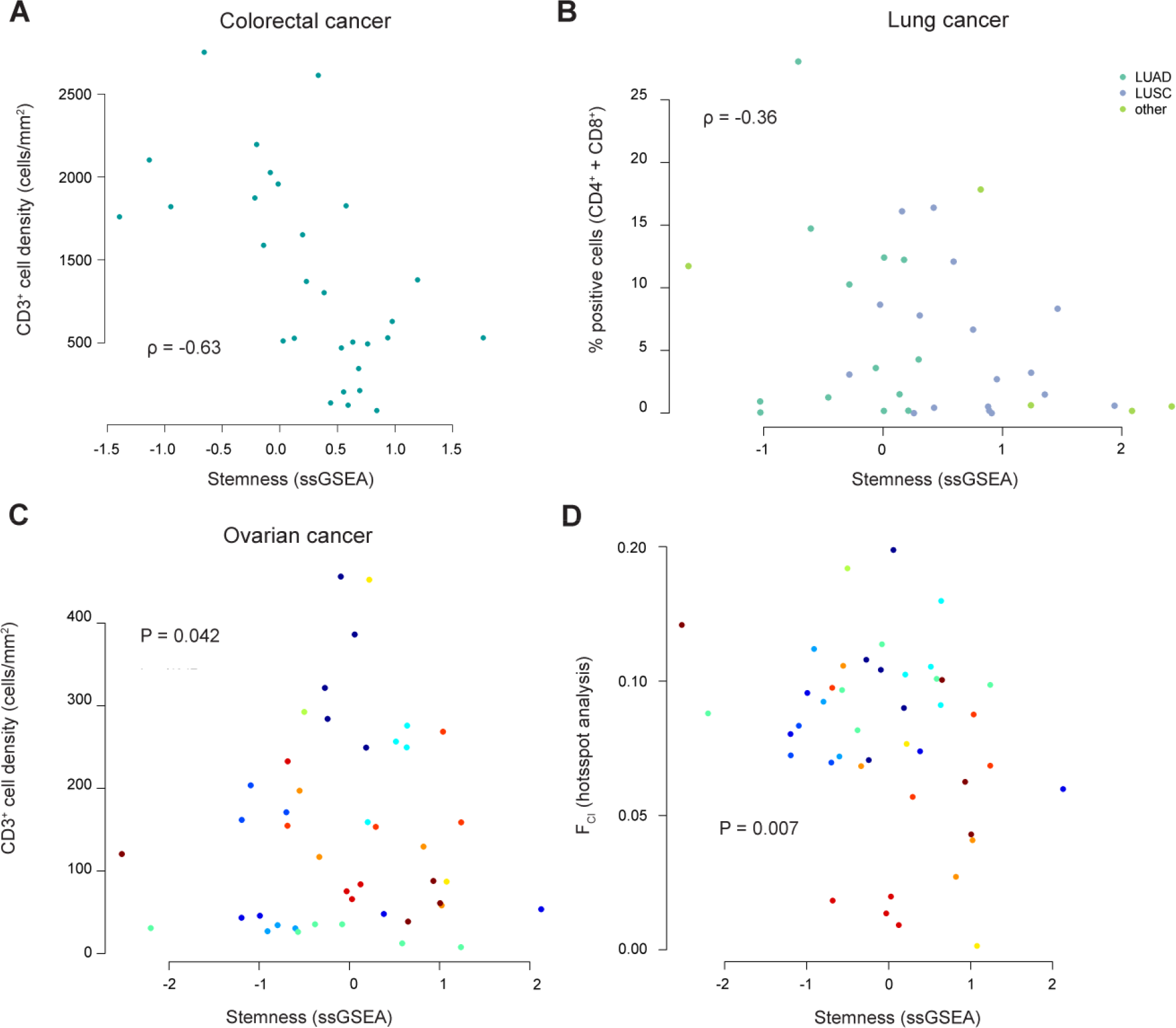
Stemness-immune signature relationship in different cancer cohorts scored via immunohistochemistry. **A)** Stemness score negatively associates with tumor-infiltrating CD3+ T cells in colorectal cancer (P < 0.001; N=33) (data from Becht et al., 2016). Each point represents one patient sample. **B)** Stemness negatively associates with tumor-infiltrating T cells (in this case the sum of CD4+ and CD8+ cells) in lung cancer (P= 0.035; N=35) (data from Mezheyeuski et al., 2018). Colors denote adeno versus squamous cell lung cancers. **C)** Stemness is significantly associated with tumor-infiltrating CD3+ T cells in a multi-site dataset from high grade ovarian cancer (P = 0.042; N = 44 samples from 12 patients) (data from Zhang et al., 2018). Colors represent individual patients. **D)** By hotspot analysis, the fraction of tissue area occupied by co-localizing tumor and immune cells (F_CI_) is negatively associated with stemness score in a multi-site dataset from high grade ovarian cancer (P < 0.01; 44 samples from 12 patients) (source data derived from Zhang et al., 2018). Colors represent individual patients.

We next evaluated this relationship in a cohort of 35 lung cancer patients with matched RNA-sequencing and IHC-based quantitation of immune cell infiltrates (Mezheyeuski et al., 2018). For consistency with the above analysis, we calculated the infiltration of T cells by summing previously calculated CD4+ and CD8+ cell fractions. Again, we observed a negative association between the stemness score and the percent of infiltrating T cells (ρ = −0.36; P = 0.035; **Figure 3B**). Furthermore, using the additional RNA-sequencing data in this cohort, we observed a negative association between the stemness and immune signature (N = 199 tumor samples; ρ = −0.19; P = 0.007).

Finally, we evaluated a small cohort of high-grade serous ovarian cancer (HGSC) cases (Zhang et al., 2018) for which matched IHC-based T cell counts and microarray-based gene expression data were available for multiple tumor sites within each patient (N = 44 samples from 12 patients). Consistent with the above findings, we found negative association between stemness and total CD3+ T cells (negative binomial mixed effects model; P = 0.042; **Figure 3C**). We did not, however, find an association between stemness and intratumoral heterogeneity in this cohort (P > 0.05), although the sample size for this analysis was even more limited (N = 25 samples from 6 patients with available data). Additionally, we previously subjected samples from this cohort to Getis-Ord Gi* ‘hotspot’ analysis to quantify immune cell engagement with tumor cells (Zhang et al., 2018). Intriguingly, all hotspot metrics showed clear negative association with stemness (mixed effects models; P < 0.01; 44 samples from 12 patients; **Figure 3D** (Nawaz et al., 2015); F_CI_ shown), suggesting that stemness negatively influences lymphocyte engagement with tumor cells.

### Stemness associates with intratumoral heterogeneity

Stemness has been implicated in fostering tumor clone diversity, with the replicative potential of CSCs enabling greater tumor heterogeneity (Greaves, 2013; Kreso and Dick, 2014). Our hypothesis suggests that stemness could additionally promote intratumoral heterogeneity by inhibiting immune selection against new tumor clones. These predictions have not, to our knowledge, been systematically tested within or across cancers. We compared stemness and intratumoral heterogeneity using data from two recent TCGA studies (See methods). Comparing stemness with predictions of cancer clonality from the first study (Andor et al., 2016) (N = 935 patients), we found a dramatic, positive correlation between median stemness and median number of clones in a cancer (ρ = 0.75; P = 0.008; N = 11 cancers; **Figure 4A**). Furthermore, we found a positive association between stemness and tumor clone count in a linear model controlling for cancer site, indicating that these trends are discernible both across and within cancers (P = 0.002, **Figure 4B**). We also exploited pan-cancer predictions of intratumoral heterogeneity from a second, larger TCGA-based study (Thorsson et al., 2018) and recovered a similarly striking trend (ρ = 0.64; P = 0.002; N = 6,791 samples across 21 cancers; **Figure 4C**), which was again significant across all samples when controlling for cancer site (P < 10^−15^; **Figure 4D**).

**Figure 4.**
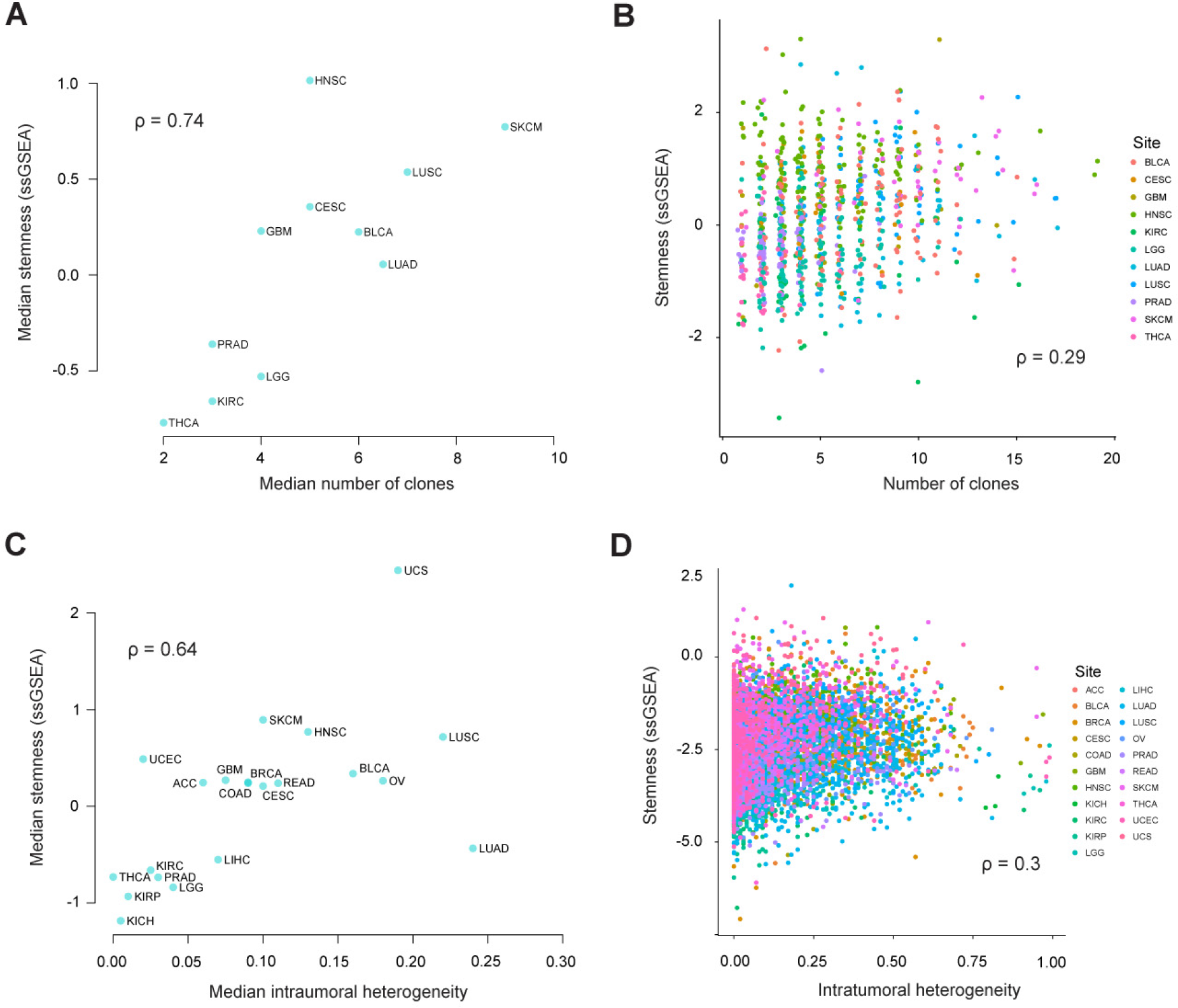
Stemness associates with intratumoral heterogeneity within and across cancers. **A)** Median stemness and median clonality, (inferred by Andor et al., 2016), are strongly correlated across cancers (N = 11; P = 0.008). **B)** Stemness score and clonality, (inferred by Andor et al., 2016), are correlated across patients while controlling for cancer type (N = 935; P = 0.008). Colored points represent different tumor sites. **C)** Median stemness score and median intratumoral heterogeneity score, (inferred by Thorsson et al., 2018), are strongly correlated across cancers (N = 20; P = 0.002). **D)** Stemness score and intratumoral heterogeneity, (inferred by Thorsson et al., 2018), are correlated across patients while controlling for cancer type (N = 6,791; P < 10^−15^). Colored points represent different tumor sites. Spearman ρ values are shown.

### Potential mechanisms and consequences of stemness-associated immunosuppression

We evaluated the association between stemness and several known genetic and environmental factors that affect antitumor immunity. Antitumor immunity involves T cell recognition of neo-antigens arising from somatic mutations (Matsushita et al., 2012). We therefore examined the association between stemness, immune signature, and non-synonymous mutation load, analyzing TCGA samples with available mutation calls (N = 6,682) (See methods). While median mutation load correlated with median immune signature across cancers (ρ = 0.49; P = 0.02, N = 21), there was generally little correlation within cancers, as has been reported in other TCGA-based analyses (**Figure S5A**, (Senbabaoglu et al., 2016)). Across cancers, median mutation load showed a positive association with median stemness (ρ = 0.61; P < 0.01, N = 21; **Figure 5A**), consistent with our observations of intratumoral heterogeneity. Indeed, we found one of the intratumoral heterogeneity estimates (of Andor et al., 2016) to be highly correlated with mutation load, although this was not the case with estimates from the more recent study (Thorsson et al., 2018). Likewise, within cancers, we generally found positive associations between mutation load and stemness (**Figure 5B**), consistent with our observation of high stemness in ultramutated endometrial cancers. We also evaluated stemness and immune associations with NetMHCpan-predicted neoantigen loads computed in the above pan-cancer analysis (Thorsson et al., 2018) and recovered qualitatively similar but slightly stronger trends (**Figure S5B and S5C**). These findings corroborate other recent studies suggesting that point mutation/neoantigen loads are not the strongest predictors of a functional antitumor immune response when considered within a cancer type (Senbabaoglu et al., 2016), underscoring the importance of other factors such as stemness, as shown here.

**Figure 5.**
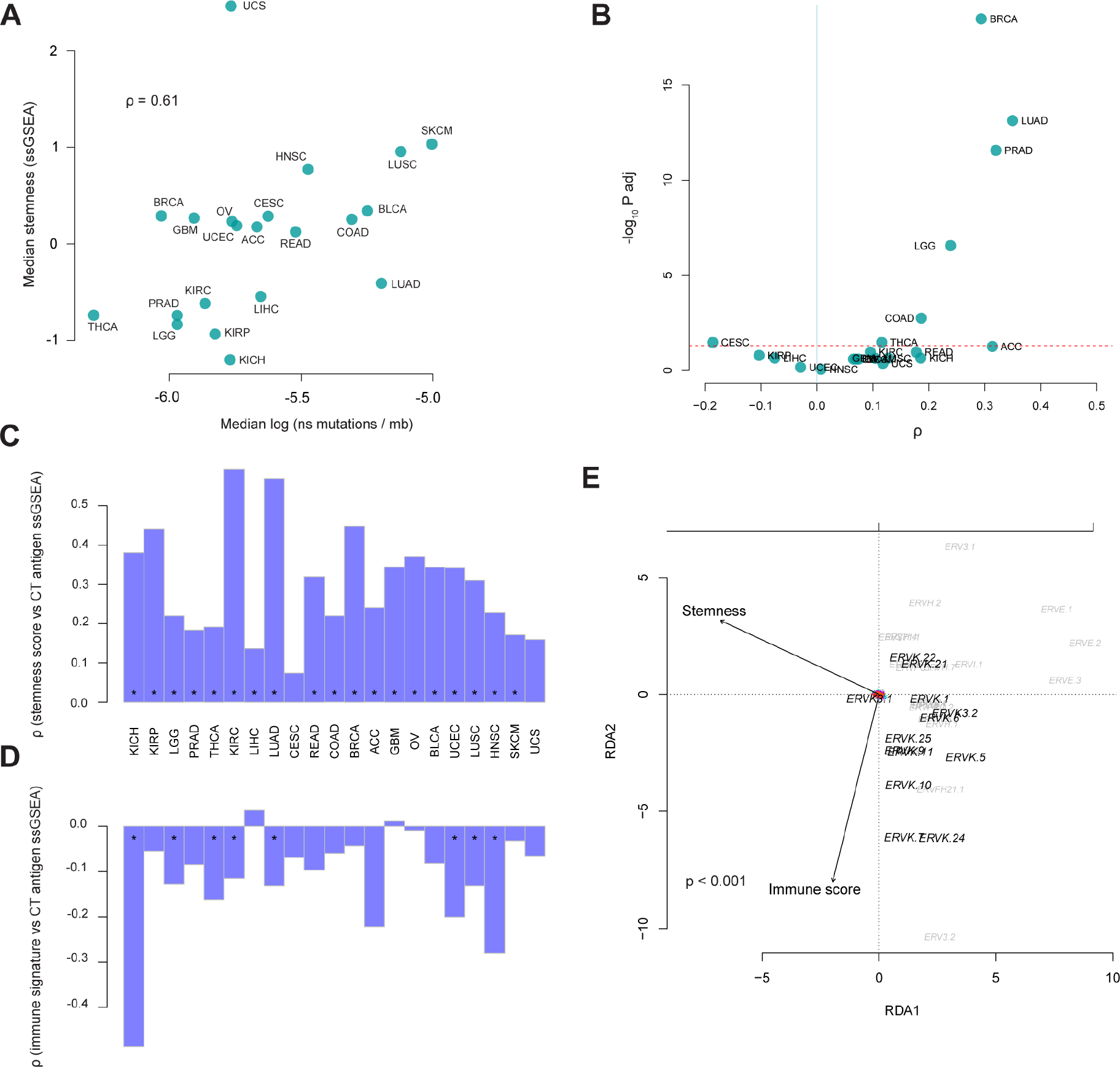
Mutation load, CT antigen expression and ERV associations with stemness. **A)** Median stemness and median mutation load are positively correlated across cancers (N = 21; P = 0.020). Mutation load is represented as log-transformed non-synonymous mutations per megabase (log ns mutations / mb). **B)** Volcano plot reveals stemness score and mutation load correlate within some cancers (upper right quadrant). The X axis represents Spearman correlation ρ values, and the Y axis represents −log 10 adjusted P values (P_adj_). Dashed red line indicates the significance threshold, P_adj_ value = 0.05. **C)** Stemness score and CT antigen expression (ssGSEA of CT antigen gene set) positively correlate in most cancers, while **D)** immune signature and CT antigen expression negatively correlate, where significant. In C and D, bar plots show the Spearman p values for each cancer type, and asterisks denote P_adj_ < 0.05. **E)** Redundancy analysis triplot reveals stemness negatively associates with multivariate ERV expression (P < 0.001; 33 ERVs evaluated in 4,252 samples, analysis conditioned by cancer type). See also **Figure S5** and **Table S2**.

Cancer-testis (CT) antigens are a class of tumor antigens that normally are expressed only in gametogenic tissue but become aberrantly expressed in a broad range of malignancies, often leading to the induction of immune responses (Simpson et al., 2005). Using a set of 201 CT genes curated by the CTdatabase (**Table S2**) (Almeida et al., 2009), we generated an ssGSEA CT antigen score and found a strong positive association with the stemness signature within cancers (**Figure 5C**). Accordingly, there was a generally negative association between CT antigen score and the immune signature, which reached statistical significance in 8 of 21 cancers (P_adj_ < 0.05; **Figure 5D**). Thus, like neoantigens, CT antigens show a positive association with stemness and a negative association with immune signature.

Normal stem cells have been shown to suppress endogenous retrovirus (ERV) expression, presumably to prevent insertional mutagenesis in long-lived stem cell lineages (Gaudet et al., 2004). Conversely, ERV expression can be activated in cancer cells (Strissel et al., 2012; Wang-Johanning et al., 2007), where it can potentially elicit antitumor immune responses by activating viral defense mechanisms and the type I interferon response (Chiappinelli et al., 2015) or by yielding immunogenic foreign epitopes (Boller et al., 1997). Despite these possibilities, clear associations between immunity and ERV expression were not seen in a recent pan-cancer analysis (Rooney et al., 2015). To better understand this relationship, we investigated interactions between stemness, immune signature, and ERV expression. Because of the repetitive nature of ERVs, we used ERV-specific read-mappings (Rooney et al., 2015) to evaluate ERV expression in 4,252 TCGA samples that overlapped with our stemness and immune signature analysis. Using redundancy analysis (a constrained extension of principal components analysis), we found that multivariate ERV expression was not clearly associated with the immune signature, consistent with prior reports (Rooney et al., 2015) (See methods); members of the ERVK family were an exception, showing moderate positive associations with immune signature (**Figure 5E**). In contrast, ERV expression showed a pervasive negative association with stemness (P < 0.001; **Figure 5E**), consistent with the notion that suppression of ERV expression is a feature of the stem cell phenotype (Yang et al., 2015). Thus, both immune signature and ERV expression are negatively associated with stemness, but these appear to be largely orthogonal relationships.

### Tumor cell intrinsic mechanisms of stemness-mediated immunosuppression

To address whether the negative association between stemness and immune signature is attributable to cancer cell-intrinsic processes, we calculated stemness scores for 1,048 cancer cell lines using gene expression data from the Cancer Cell Line Encyclopedia (CCLE) (Barretina et al., 2012) (See methods). We first assessed associations between stemness scores and the expression of 11 mapped ERVs in CCLE transcriptomes and found that 3/3 ERVs with non-negligible expression levels across cell lines were significantly negatively associated with stemness, two of which remained significant when controlling for tissue of origin (linear models; P_adj_ < 0.05; **Figure S6)**. We next generated an ssGSEA score for type I interferon signaling (Reactome IFN alpha/beta pathway gene set) and observed a clear negative relationship between this signature and stemness (ρ = −0.22. P < 10^−10^) (**Figure 6A);** this association remained significant when controlling for tissue of origin of the cell line (linear model; P < 0.001) and when omitting cell lines derived from hematopoietic lineages (linear model; P = 0.02). To evaluate the association between stemness and type I IFN signaling in non-neoplastic lineages, we took advantage of the stem cell gene expression dataset previously used to validate stemness signatures (GSE30652). This too revealed a striking negative association between type I IFN signaling and stemness (ρ = −0.81; P < 10^−15^; **Figure 6B**).

**Figure 6:**
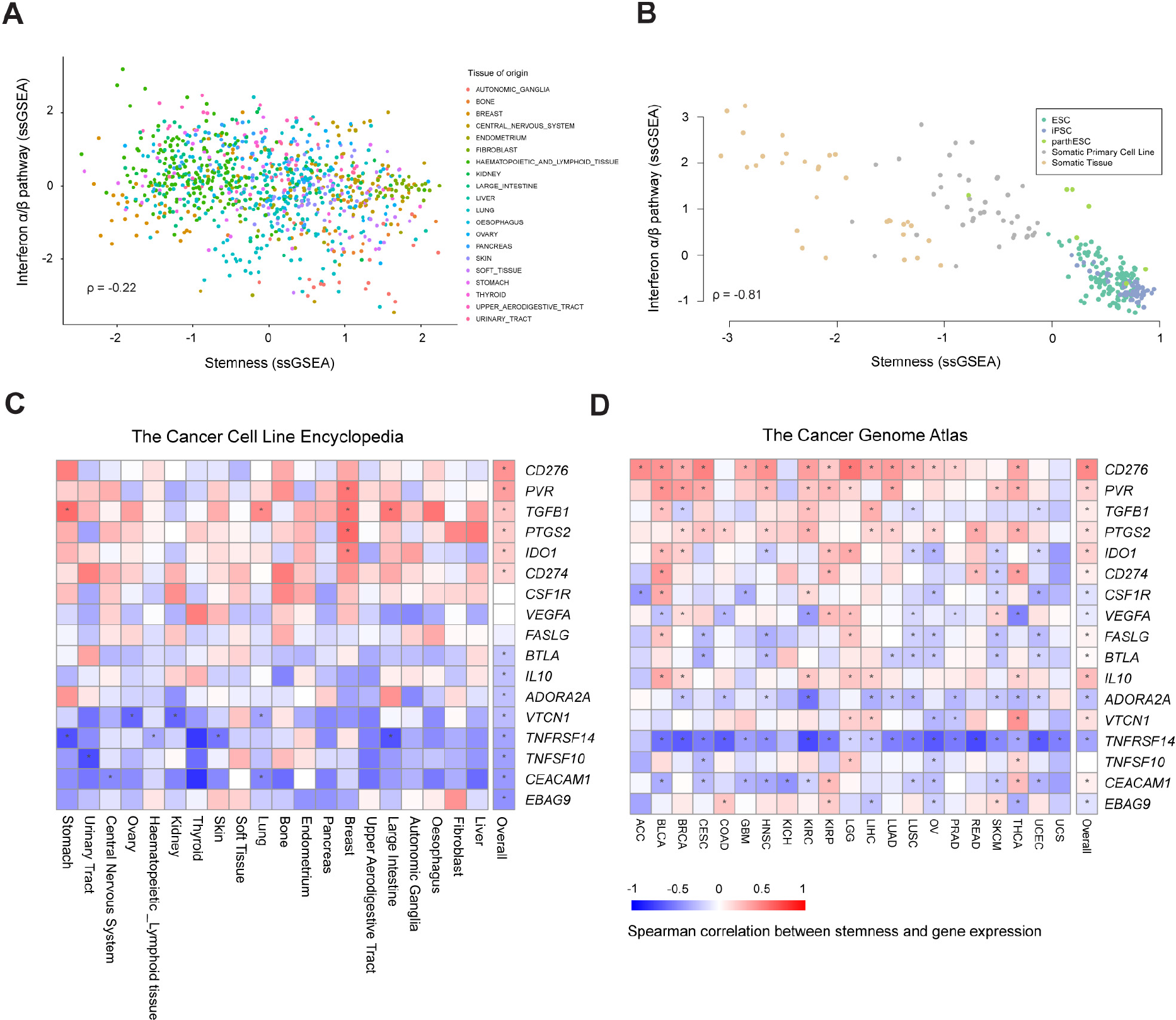
Cell-intrinsic stemness score associate with decreased type I interferon signaling and increased expression of *CD276* and *PVR*. In A and B, colored points represent different cell lines and tissues in data from the CCLE and GSE30652, respectively. **A)** Stemness score negatively associates with type I interferon signaling (P < 10^−10^; Reactome IFN α/β pathway ssGSEA) across cancer cell lines. Only tissues represented by more than 10 cells lines were included in analysis. **B)** Stemness negatively associates with type I interferon signaling across non-malignant stem cells, somatic tissues, and primary cells (P < 10^−15^). **C, D)** Heatmaps showing Spearman correlations for stemness and select immunoinhibitory genes based on data from the CCLE (C) and TCGA (D). Spearman correlations were calculated within tissues represented in the CCLE by more than 10 cells lines. Genes are ranked according to the final column (“overall”), which represents the correlation across all samples, irrespective of cell line or tumor type. Red-blue intensities reflect the correlation ρ values. Asterisks denote Benjamini-Hochberg-corrected significant associations (P_adj_ < 0.05). See also **Figure S6**.

Finally, to examine other cell-intrinsic mechanisms of immunosuppression, we analyzed a curated list of immunoinhibitory genes previously reported to be expressed in human cancer cells. Using both the CCLE and pan-cancer TCGA datasets, the expression of each of these genes was assessed in relation to our stemness signature (**Figures 6C and 6D**). This revealed positive associations between stemness and a number of immunoinhibitory genes, including *CD276* (*B7-H3*, shown to inhibit T cell activation and autoimmunity (Lee et al., 2017), *PVR* (*CD155*, a member of the B7/CD28 superfamily, shown to exhibit potent inhibitory action in different subsets of immune cells (Mahnke and Enk, 2016), and *TGFB1* (a key player in the induction of immunological tolerance (Johnston et al., 2016). Thus, the stemness phenotype involves the expression of several gene products that could potentially serve as targets for immune modulation.

## Discussion

Although cancer stemness, antitumor immunity, and intratumoral heterogeneity have all emerged as centrally important features of cancer in recent years, their covariation across cancers has not been systematically investigated. Here we report that stemness is associated with suppressed immune response, higher intratumoral heterogeneity, and dramatically worse outcome for many cancers. Although correlative analyses such as ours do not reveal causality, we propose that the stemness phenotype found in cancer cells, similar to that in normal stem cells, involves the expression of immunosuppressive factors that engender the formation of immune-privileged microenvironments in which tumor clone diversification can occur. The resulting heterogeneity provides a substrate for the selection of treatment-resistant tumor clones, resulting in inferior clinical outcomes. Thus, our findings implicate the stemness phenotype as a shared therapeutic target to achieve the dual objectives of constraining tumor evolution and enhancing antitumor immunity.

A recent pan-cancer analysis reported inconsistent relationships between cancer stemness and immunity, recovering negative relationships between stemness and tumor-infiltrating lymphocytes for some cancers and positive relationships for others (Malta et al., 2018). While this work provided a valuable perspective on stemness across cancers, our results differ in that we recover much stronger and more pervasive negative relationships with immune infiltration and survival. In contrast to ssGSEA, the OCLR approach used above was trained on a cohort that lacked any malignant samples. Moreover, when we reproduced their analysis, we found many components of their stemness score (i.e. positive model weights) were immunologically relevant genes; for example, among the 50 most positive gene weights were *IDO1, LCK, KLRG2, PSMB9* (a component of the immunoproteasome), and multiple TNF-receptors. With the OCLR approach, higher expression of such immune genes contributes positively to the stemness score, precluding an unbiased assessment of the relationship between stemness and tumor immunity.

The contribution of cancer stemness to intratumoral heterogeneity has been postulated for some time (Kreso and Dick, 2014), but direct evidence has been lacking. We recovered a dramatic positive association between stemness and intratumoral heterogeneity across cancers, which is especially noteworthy given that these metrics were derived from separate data types (i.e. mRNA vs DNA). Given recent work from our group and others linking increased intratumoral heterogeneity with decreased immune cell infiltration (Safonov et al., 2017; Zhang et al., 2018), one could speculate that stemness might enable intratumoral heterogeneity by both increasing the replicative capacities of individual tumor clones and by shielding antigenic clones from elimination by the immune system.

We found generally positive associations between stemness and mutation load within cancers, and clear evidence of this across cancers, consistent with studies demonstrating accumulation of mutations in normal adult stem cells (Blokzijl et al., 2016). We also found strong positive associations between stemness and CT antigen expression, which is consistent with reports of CT antigen expression in mesenchymal (Saldanha-Araujo et al., 2010) and embryonic stem cells (Lifantseva et al., 2011), as well as cancer stem cells (Yamada et al., 2013). It is also consistent with prior studies (Thorsson et al., 2018) and the present work finding non-significant or negative correlations between immune infiltration and CT antigen expression (**Figure 5D**). Thus, the negative association between stemness and immunity is not attributable to low neoantigen or CT antigen load, strongly implicating the involvement of other mechanisms.

ERVs, which constitute approximately 8% of the human genome, are known to be suppressed in pluripotent and embryonic stem cells (Yang et al., 2015) yet activated in human cancer (Roulois et al., 2015; Schmitt et al., 2013), leading us to ask which behavior would predominate in high stemness cancers. We found a strong negative association between stemness and ERV expression, indicating that the stemness phenotype in human cancer retains this property of normal stem cells. This negative association was also seen in cancer cell line data, suggesting a cell-intrinsic phenomenon. The cell line data also revealed a negative correlation between stemness and type I IFN signaling. Whether ERV suppression underlies the low intrinsic IFN signaling in high stemness cancer cell lines awaits experimental investigation. However, in support of this notion, an attenuated innate immune response is a major characteristic of embryonic stem cells (Guo et al., 2015), and we find clear confirmation of this in non-neoplastic stem cell transcriptomes (**Figure 6B**). Collectively, these data suggest that activation of a stemness program in tumors could limit antitumor immune responses by silencing ERVs and repressing type I IFN signaling in a cell-intrinsic manner (Fuertes et al., 2013).

Using CCLE data, we found a clear association between stemness and the expression of several immunoinhibitory genes, including *CD276, PVR*, and *TGFB1*. These associations are especially intriguing given that CCLE-based gene expression profiles are independent of any ongoing influence of the immune system. CD276, a B7 family ligand, is now being clinically targeted due to expression on both cancer cells and tumor infiltrating blood vessels (Seaman et al., 2017). Intriguingly, CD276 is co-expressed with CD133, a marker that distinguishes cell populations enriched for cancer stem cells in colorectal cancer (Bin et al., 2014). PVR is a key ligand in an emerging checkpoint pathway involving TIGIT, an inhibitory receptor expressed on T cells and other immune cells (Mahnke and Enk, 2016). Although no association between stem cells and PVR has been described so far in mammalian stem cells, expression of PVR can be activated by sonic hedgehog signaling (Solecki et al., 2002), a pathway essential for self-renewal and cell fate determination in normal and cancer stem cells (Cochrane et al., 2015). TGFB1 and other TGFB family members have well documented roles in development (Watabe and Miyazono, 2009) and CSC proliferation and maintenance (Ikushima et al., 2009; Penuelas et al., 2009).

Recent work has demonstrated that tumor-intrinsic oncogenic signaling pathways have immune suppressive properties (Spranger and Gajewski, 2016, 2018), but this too can be understood in the context of stemness. For example, molecular pathways involving WNT/B-catenin, MYC, PTEN and LKB1 have been implicated in the inhibition of antitumor immunity (Spranger and Gajewski, 2018), yet they also play important roles in stem cell maintenance (Gurumurthy et al., 2010; Hill and Wu, 2009; Murphy et al., 2005; Nusse, 2008). Thus, stemness may provide a unifying framework for understanding how various oncogenic signaling pathways engender an immunosuppressive tumor microenvironment.

Tumors with reduced lymphocyte infiltration are less susceptible to immunotherapies, including checkpoint blockade (Ji et al., 2012). Consequently, our findings imply that high-stemness tumors will be more refractory to immunotherapy, much as they are to more conventional chemo- and radiotherapies (Morrison et al., 2012). If stemness is an underlying cause of the cold tumor phenotype, it may prove beneficial to target specific molecules or pathways that appear to be inherent to the stemness phenotype, such as the aforementioned immunoinhibitory molecules. Likewise, high-stemness cancer cells might be rendered more sensitive to immunotherapy by administering drugs that induce cell differentiation to irreversibly disrupt the stemness phenotype (de The, 2018). This would bring the additional benefit of constraining further tumor evolution, creating the conditions for durable clinical responses.

## Methods

### 1 Contact for reagent and resource sharing

Further information and requests for resources should be directed to and will be fulfilled by the lead contact, Brad Nelson.

### 2 Experimental models and subject details

Clinical parameters and molecular subtypes for TCGA data were extracted from the expression data files accessed through *TCGAbiolinks* package. Mutation data were downloaded from Firebrowse (www.firebrowse.org) as the number of non-synonymous mutations. Tumor purity (Aran et al., 2015), mRNAsi (Malta et al., 2018), intratumoral heterogeneity (Andor et al., 2016; Thorsson et al., 2018), and CIBERSORT and neoantigen scores (Thorsson et al., 2018) by extracting the relevant data for overlapping samples from the respective supplemental materials.

For correlations between stemness scores and IHC-based immune cell counts, immune cell infiltration data were provided by the authors of the respective studies (Becht et al., 2016; Mezheyeuski et al., 2018). Overlapping expression data (microarray or RNA-sequencing) was obtained from the gene expression omnibus (GEO: GSE39582; GEO: GSE81089), or provided by the authors (Bashashati et al., 2013; Zhang et al., 2018).

### 3 Method Details

All analyses were conducted with R v. 3.4.2. We accessed RNA sequencing as upper quartile normalized FPKM using the the *TCGAbiolinks* R/Bioconductor package (Colaprico et al., 2016), for each cancer of interest, and expression data were merged across cancers. For genes with multiple annotated transcripts, we selected the transcript with the greatest expression to represent the gene, then filtered the expression set to include only primary samples (except for melanoma, for which we included metastases), removed patients with duplicate samples, and removed any patients without a consensus purity score in (Aran et al., 2015) to enable purity corrections in analyses. For microarray datasets, we converted probe IDs to human gene symbols using *biomaRt* (Durinck et al., 2009), and retained the most expressed probe for each gene, as above. Where appropriate (e.g. for linear modelling), expression data were log_2_(x + 1) transformed.

We calculated stemness and other ssGSEA signatures using the *GSVA* package in R (gene sets in table S1; (Hanzelmann et al., 2013)) without normalization, and subsequently scaled values as z-scores within the dataset of interest. *xCell* enrichment scores were calculated in R, using the *rawEnrichmentanalysis* (Aran et al., 2017) function, which omits scaling scores to [0, 1] and correction for correlations among related cell types (Aran et al., 2017), as we sought to avoid introducing non-linearities from these steps into analysis. To generate the immune signature, we summed z-scored signatures of cell types of interest (CD8+ T cells, NK cells, B cells). Because z-scoring of ssGSEA scores was done within each dataset, these scores should not be directly compared across datasets.

### 4 Quantification and Statistical analysis

We used non-parametric Spearman’s correlation to assess pairwise associations between variables of interest within cancers, or for median values across cancers, adjusting for multiple tests using the Benjamini-Hochberg method, where appropriate. For analyses across multiple cancer types or subtypes, we used linear models, controlling for site or subtype as fixed main effects, and inspecting model residuals to ensure model assumptions were reasonable. For analyses controlling for purity, purity (as consensus purity estimate from (Aran et al., 2015)) was included as a covariate in linear models.

We conducted survival analyses using Cox proportional hazards models, calculating 95% CIs on log hazard ratios. We tested model assumptions using *cox.zph* (Therneau, 2018). Where there were significant violations of model assumptions (P < 0.05), we inspected model Schoenfield residuals; we found that for the few instances in which model assumptions were violated, this was attributable to higher than predicted survival for some long-term survivors. For analyses across all patients and cancer types, we stratified Cox models by cancer type. For IHC data with multiple samples taken from the same patient, we modelled patient as a random intercept in mixed effects models implemented in *lme4* in R (Bates et al., 2015) and assessed the significance of fixed effects using likelihood-ratio tests on nested models.

Pathway enrichment analysis was conducted using *ReactomePA* (Yu and He, 2016) after testing for differential expression using *limma* (Ritchie et al., 2015). For enrichment analysis, we selected the top 1000 significantly down-regulated genes in high stemness cancers based on the moderated t statistic; in cancers where < 1000 genes were significantly down-regulated (P_adj_ < 0.05), we selected all significantly down-regulated genes for downstream analysis. Significance of enrichment was evaluated at P_adj_ < 0.05, and recurrently enriched pathways were defined as those that were significantly enriched in the greatest number of cancers. In *limma* analyses that included purity as a covariate, purity was log-transformed for consistency with the transformation of expression values.

To assess ERV expression, we first variance-filtered mapped ERVs to select those above the median interquartile range of expression using the *genefilter* R package (R. Gentleman et al., 2018). We conducted partial redundancy analysis using the *vegan* package, implementing the default distance metric (Euclidean). We conditioned the analysis by cancer type to control for cancer type-specific effects, and tested the significance of multivariate associations using permutation tests (n=1000) in *vegan* (J. et al., 2015).

### 5 Data and software availability

All data used in this analysis are publically available (see Model and Subject Details) or provided by the original authors. Scripts to reproduce analyses are available upon request.

## Acknowledgments

Funding was provided by the BC Cancer Foundation (B.H.N), Canadian Cancer Society Research Institute (B.H.N), Canada’s Networks and Centres of Excellence (B.H.N), Canadian Institutes of Health Research (B.H.N), Terry Fox Research Institute (B.H.N.), Cancer Research Society (B.H.N.), Vanier Canada Graduate Scholarship (A.W.Z) and Canadian Institutes of Health Research Postdoctoral Fellowships (P.T.H). The results shown here are in whole or in part based upon data generated by the TCGA Research Network https://cancergenome.nih.gov/ or the CCLE collection https://portals.broadinstitute.org/ccle.

We thank Dr. John Dick (University of Toronto) and Dr. Julian Lum (BC Cancer) for critical review of the manuscript and Dr. Swetansu Pattnaik (Garvan Institute of Medical Research) and Dr. Peter Watson (BC Cancer) for helpful discussions.

## Author Contributions

Study design, A.M., P.T.H., B.H.N.; Data analysis and interpretation, P.T.H., A.M., A.W.Z, B. H.N.; Immunohistochemistry, J.B., P.M., A.M., E.B., A.D.R.; Manuscript writing, A.M., P.T.H., B.H.N.; Manuscript review, A.M., P.T.H., J.S, A.W.Z, A.M, E.B, A.D.R, B.H.N.; Study supervision, B.H.N.

